# The retina’s neurovascular unit: Müller glial sheaths and neuronal contacts

**DOI:** 10.1101/2024.04.30.591885

**Authors:** William N. Grimes, David M. Berson, Adit Sabnis, Mrinalini Hoon, Raunak Sinha, Hua Tian, Jeffrey S. Diamond

**Author notes:** equal contributions (William N. Grimes, David M. Berson, Adit Sabnis).

## Abstract

The neurovascular unit (NVU), comprising vascular, glial and neural elements, supports the energetic demands of neural computation, but this aspect of the retina’s trilaminar vessel network is poorly understood. Only the innermost vessel layer – the superficial vascular plexus (SVP) – is ensheathed by astrocytes, like brain capillaries, whereas glial ensheathment in other layers derives from radial Müller glia. Using serial electron microscopy reconstructions from mouse and primate retina, we find that Müller processes cover capillaries in a tessellating pattern, mirroring the tiled astrocytic endfeet wrapping brain capillaries. However, gaps in the Müller sheath, found mainly in the intermediate vascular plexus (IVP), permit different neuron types to contact pericytes and the endothelial cells directly. Pericyte somata are a favored target, often at spine-like structures with a reduced or absent vascular basement lamina. Focal application of adenosine triphosphate (ATP) to the vitreal surface evoked Ca^2+^ signals in Müller sheaths in all three vascular layers. Pharmacological experiments confirmed that Müller sheaths express purinergic receptors that, when activated, trigger intracellular Ca^2+^ signals that are amplified by IP_3_-controlled intracellular Ca^2+^ stores. When rod photoreceptors die in a mouse model of retinitis pigmentosa (*rd10*), Müller sheaths dissociate from the deep vascular plexus (DVP) but are largely unchanged within the IVP or SVP. Thus, Müller glia interact with retinal vessels in a laminar, compartmentalized manner: glial sheathes are virtually complete in the SVP but fenestrated in the IVP, permitting direct neural-to-vascular contacts. In the DVP, the glial sheath is only modestly fenestrated and is vulnerable to photoreceptor degeneration.

## Introduction

Retinal processing, like all neural computation, is metabolically expensive and requires extensive vascular support. Throughout the brain, pericytes, endothelial cells and perivascular glia cooperate to 1) extract metabolic resources from the blood; 2) remove waste; 3) maintain the blood brain barrier (BBB); and 4) dynamically control blood flow in response to changes in neural activity, a process known as neurovascular coupling (NVC) or functional hyperemia.

Endothelial cells forming the capillary wall make tight junctions with one another, an essential element of the BBB. Endothelial cells are coated in basement lamina and partially covered by pericytes, contractile cells that cling to vessels’ external surface and are also surrounded by basement lamina (Attwell et al., 2016; Kovacs-Oller et al., 2020).

Most mammals, including humans, have highly vascularized retinas (Figure 1). Even brief interruptions in blood flow dramatically diminish retinal light responses (for review see (Osborne et al., 2004)). While photoreceptors are nourished by the choroid, all other retinal neurons rely on the central retinal artery for their blood supply. This artery branches upon entering the eye to form a trilaminar vascular network comprising the superficial, intermediate, and deep vascular plexuses (SVP, IVP, and DVP; Figure 1). The SVP lies within the ganglion cell layer (GCL) and optic fiber layer and includes veins, arterioles, and capillaries (Kornfield & Newman, 2014; Snodderly et al., 1992). The IVP and DVP consist mainly of capillaries. The IVP is situated at the boundary between the inner nuclear layer (INL) and the inner plexiform layer (IPL), while the DVP lies in the outer plexiform layer (OPL) near the photoreceptor synaptic terminals.

**Figure 1.**
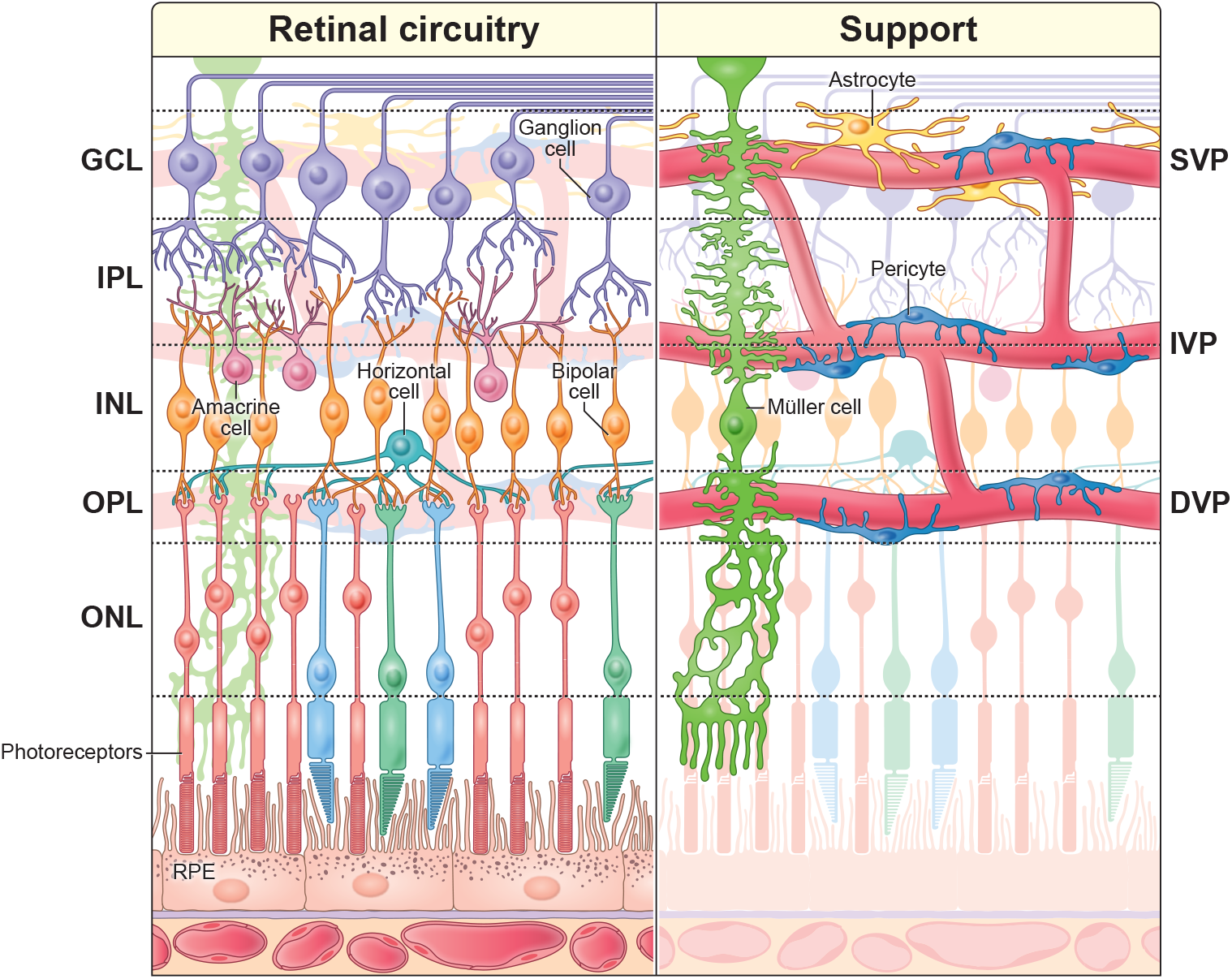
Organization of the mammalian retina. *Left* The retina is a laminar structure comprising five neuron classes: photoreceptors, horizontal cells, bipolar cells, amacrine cells, and ganglion cells. *Right*, Retinal circuitry is supported by a network of blood vessels, pericytes, and glial cells. Capillaries are wrapped by contractile pericytes, which regulate blood flow. Retinal astrocytes are restricted to the SVP (see also Figure S1). Radial Müller glia extend through all layers of the retinal circuitry, interacting with both capillaries and neurons. GCL = ganglion cell layer, IPL = inner plexiform layer, INL = inner nuclear layer, OPL = outer plexiform layer, ONL = outer nuclear layer, RPE = retinal pigment epithelium, IVP = intermediate vascular plexus, DVP = deep vascular plexus.

In the central nervous system, capillaries and pericytes are ensheathed by glial endfeet. In the brain, the endfeet derive from astrocytes (Mathiisen et al., 2010), but in the retina astrocytes are confined to the SVP, where they, together with radial Müller glia, wrap vessels (Albargothy et al., 2023) (Figure 1, S1). In the IVP and DVP, only Müller glia wrap capillaries, as shown in cat (Hollander et al., 1991), tree shrew (Ochs et al., 2000; Reichenbach et al., 1995), and human (Erskine, 1963). Müller cells span nearly the entire depth of the retina, extending fine processes laterally in all synaptic and vascular layers (Figure 1, 2A-D).

**Figure 2.**
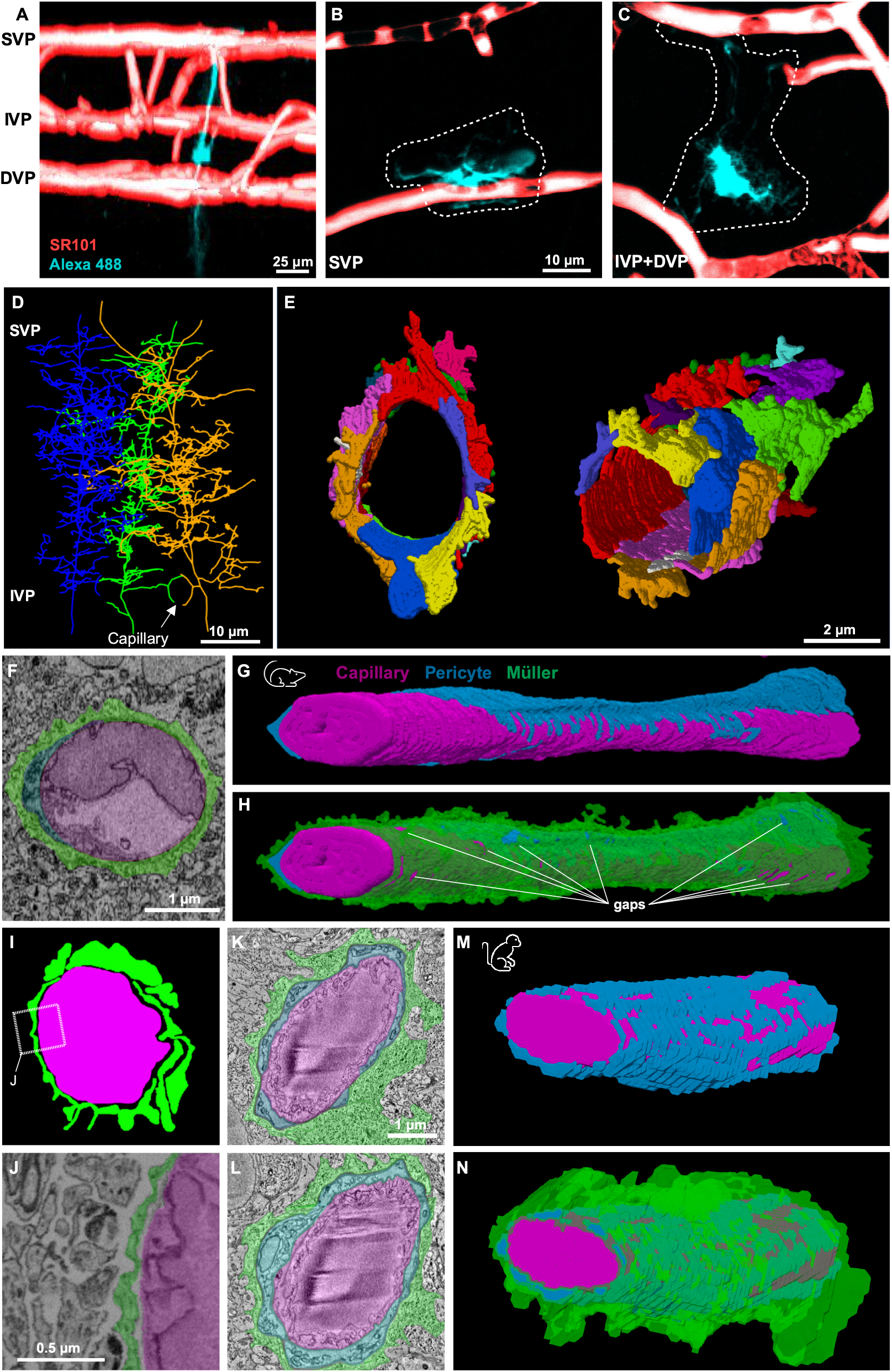
Müller processes form sheaths around pericytes and capillaries in mouse and primate retinas. A-C) 3D rendering of a Müller glia (teal, injected with Alexa 488) and the vasculature (red, SR-101 pretreatment). B-C) Z-projections of the superficial (B) and the intermediate + deep (C) layers. D-E) Müller ultrastructure in the IPL revealed by SBFSEM (Ding et al., 2016). D) Three partially reconstructed Müller glia. A capillary (not shown) passes through a ring formed by neighboring Müller cells. E) Volumetric reconstruction of glial processes surrounding a IVP capillary. Different Müller processes are colored distinctly to indicate the tessellated coverage. F) Electron micrograph of a retinal capillary (*magenta*) in cross section highlighting the anatomical arrangement with pericytes (*blue*) and Müller glia (*green*). G) Volumetric reconstruction of an IVP capillary and the surrounding pericytes. H) Same as in G with the addition of the Müller sheath. I) Colored image of a retinal capillary taken from an EM block in which extracellular space has been preserved (Palatto et al., 2016). J) Zoomed in electron micrograph (from I) shows extracellular space between neurons and nearly-complete Müller sheath. K-N) Ultrastructure of a retinal capillary from the IVP in a macaque (same color scheme as for other panels), similar to the arrangement found in mouse. K,L) Capillary cross sections. M,N) 3D reconstructions showing the IVP capillary and its wrapping by pericyte and Müller processes (I). See also Figure S2.

Visual stimulation dilates blood vessels in all three vascular layers (Kornfield & Newman, 2014). The IVP is particularly interesting in this context: In healthy animals, oxygen tension is lowest near the IVP (Yu et al., 1994), indicating high oxygen consumption relative to replenishment. The IVP also exhibits the largest relative changes in capillary diameter in response to visual stimulation (Kornfield & Newman, 2014), and functional studies showed that local Ca^2+^ signals in Müller cells are linked to IVP capillary dilation (Biesecker et al., 2016), suggesting that the architecture of the IVP NVU might differ from that in the SVP and DVP.

We therefore undertook the first large-scale ultrastructural reconstructions of glial and vascular cells in the IVP and DVP and their Müller cell partners using serial blockface scanning electron microscopy (SBFSEM). We found that Müller glia collectively provide near-complete ensheathment of retinal capillaries in mice and non-human primates (>90% coverage).

Nonetheless, gaps in the sheath, mostly in the IVP, allow pericytes and endothelial cells to make direct contact with neurons, including bipolar, horizontal, amacrine, and ganglion cells. Pericyte somas are a favored target of neuronal contacts, often at complex spine-like appendages where the basement lamina is thinned or absent. Similar neurovascular architecture, including fenestrations in the glial sheaths permitting neuronal contact onto pericytes and endothelial cells are apparent in SBFEM datasets of the neocortex.

In live imaging experiments, exogenously applied ATP elicited Ca^2+^ signals throughout Müller cells, including the sheaths covering all three vascular layers. Pharmacological experiments suggested that Ca^2+^ signals reflect activation of Müller cell purinergic receptors driving calcium mobilization via inositol 1,4,5-trisphosphate receptors (IP_3_Rs). In the early stages of retinal degeneration in the *rd10* mouse model (Ivanova et al., 2019), we found Müller sheaths were altered mostly in the DVP, near the photoreceptors.

Taken together, these findings provide new insights into the structural organization of the neurovascular unit in the mammalian retina. They identify novel interaction sites between neurons, pericytes and endothelial cells and demonstrate that the NVU is disrupted in a layer-specific manner in early stages of retinal degeneration.

## Results

### Nearly complete ensheathment of retinal vessels by Müller glia

We examined publicly available SBFSEM datasets of C57/Bl6 mouse retina (Ding et al., 2016; Helmstaedter et al., 2013; Pallotto et al., 2015) to visualize glial ensheathment of retinal vessels at unprecedented resolution and scale. Our study builds upon an earlier SEM analysis in mouse retina focused on astrocytic sheathes in the SVP (Albargothy et al., 2023). We focused mainly on the IVP and DVP, where Müller cells rather than astrocytes wrap the capillaries. We also examined a new small-volume 3D EM dataset of macaque IPL (unpublished dataset, see Star Methods).

We started with the Ding dataset, the largest of the mouse retinal datasets (>200×200 µm^2^) that contains the entire IVP, although SVP vessels are mostly cropped, and DVP vessels are excluded completely. Müller cells were easily recognizable even in single sections; most fine distal processes exhibit distinctively concave surfaces as they fill the spaces between the largely convex neuronal profiles. They also exhibit distinctive patterning of intracellular organelles.

We generated skeleton reconstructions of several neighboring Müller cells (Figure 2D) with single vertical stalks spanning the IPL and profuse, highly branched fine processes extending laterally in the IPL, often wrapping nearby IVP capillaries (Figure 2D *arrow*).

We made volume reconstructions of the Müller processes one IVP capillary (Figure 2E).

Numerous glial endfeet tessellated at the vessel surface to form a sleeve that was locally continuous. This quilt of endfeet could be traced back to just four Müller cells (Figure S2). This may reflect specific involvement of certain Müller cells, because >12 different Müller are close enough to contribute to glial sheaths at any point in the vascular plexus. That inference is based on inter-stalk spacing of ∼7 µm in a roughly hexagonal array and ∼10 µm of lateral extent of Müller cell processes at the IVP level (Figure 2D). More extensive reconstructions are required to determine the degree of convergence among Müller cells in various vascular compartments. No gaps in coverage were apparent in the stretch of capillary in Figure 2E, but more extensive reconstructions revealed sparse gaps (Figure 2G,H) that are explored in detail below.

We analyzed a short segment (∼5 µm) of IVP capillary in a second mouse retina SBFSEM dataset in which fixation enhanced the extracellular space to highlight points of contact and adhesion between cells (Pallotto et al., 2015). A clear gap separated the Müller processes from the vascular wall, but the Müller processes themselves adhered to one another, forming a continuous glial sheath (Figure 2I-J). This suggests that the extensive Müller ensheathment observed in the Ding dataset is not an artifact of glial swelling that can occur during chemical fixation (Korogod et al., 2015).

In the macaque retinal SBFSEM volume, a reconstructed ∼10 µm extent of IVP capillary revealed virtually complete coverage by Müller endfeet, wrapping both pericytes and endothelial cells (Figure 2K-N). No gaps could be identified between glial contacts in this small volume.

Thus, in mice and primates, IVP pericytes and endothelial cells are mostly shielded from the neuropil and from membrane-impermeant signaling molecules released by retinal neurons.

### Fenestrations in Müller sheaths permit direct interactions between neurons and IVP capillaries

In the Ding volume, we systematically searched all vessels for gaps in the glial sheath where neurons contacted pericytes and endothelial cells at the vessel wall (Figure 3A-D). Because terminal pericyte processes are very fine and easily confused with endothelial processes, we made full skeleton reconstructions of all pericytes in the volume (Figure 3A,C). Pericytes tiled the vessels of the IVP with minimal overlap, as well as the vessels connecting the IVP with the SVP (Figure 3A). Neuronal contacts onto pericytes (green dots in Figure 3B) and endothelial cells (purple dots) (total n=1631) were almost entirely restricted to the IVP. Just a few contacts were scattered along the vessels connecting the SVP and IVP. Roughly half (58%) of the contacts were made onto the exposed pericyte surface, with the remainder onto exposed endothelial cells (Figure 3B,D).

**Figure 3.**
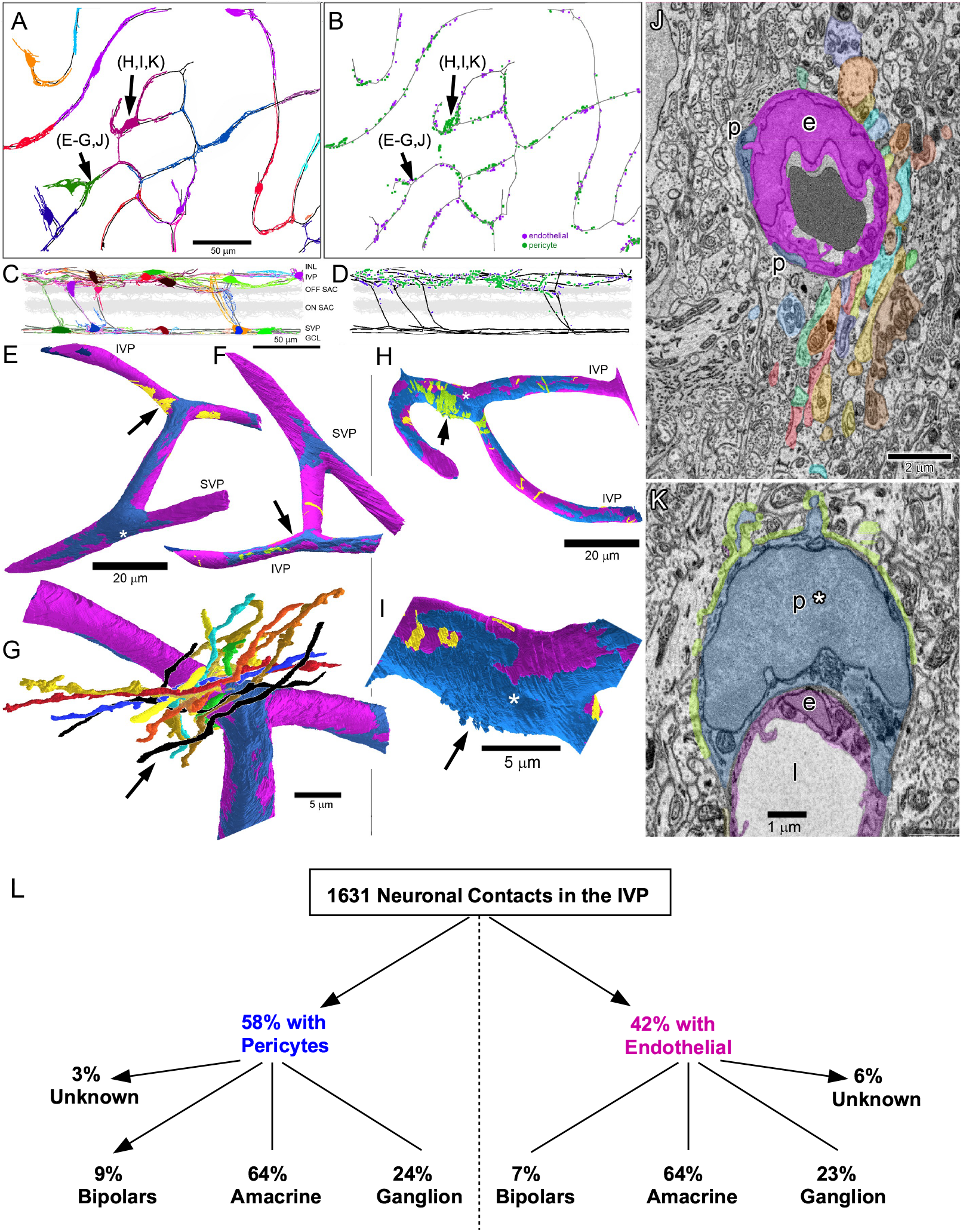
Direct contacts between neurons and vascular elements at Müller sheath gaps. A) Top-down (*en face*) views of the entire mouse SBFSEM volume (Ding et al., 2016) showing IVP capillaries (black contours) and their associated pericytes (colored skeletons). Linker vessels to the SVP are also shown (SVP vessels omitted for clarity). B) Map of neuronal contacts onto the basement lamina of either endothelial cells (*purple dots*) or pericytes (*green dots*); black vessels and scale as in A. C,D) Vertical (side) views of the data in A and B, respectively, except that the SVP and its associated pericytes have been included in C. Scale in C applies to D. E,F,G) Volume reconstructions of the region of the vascular network indicated by *black arrows* in A, B, E, F and G. Endothelium (*magenta*), pericytes (*dark blue*). E and F show neuronal contacts onto endothelial basement lamina (*yellow*) and pericyte basement lamina (*green*). The SVP pericyte soma (asterisk) is the source of all pericyte processes in E-G except for the darker blue ones (E, upper left). G shows an enlarged view of the region marked by the yellow patch and black arrow in E along with volume reconstructions of a subset of cofasciculating neuronal processes making endothelial contact. Retinal ganglion cell dendrites (*black*), amacrine processes (*other colors*). H,I) Volume reconstructions of the region of the IVP indicated by *yellow arrows* in A and B. Perspective is from the vitreal side; conventions as in E-G. Here, neuronal contacts are mainly onto pericyte (*green*) rather than endothelium (*yellow*). Contacts are concentrated on the soma of the central pericyte (asterisk), which gives rise to all pericyte processes shown except for the darker blue ones at lower right in H. Panel I shows an enlargement of the somatic region of the pericyte with the green marker of neuronal contact omitted to reveal spine-like appendages of the soma (blue arrow). J) Representative electron micrograph showing a slice through the region indicated by the arrow in G, and following the same color conventions (e: endothelial cell; p: pericyte processes). K) As in J, but drawn from the region marked by the blue arrow in H and I. Points of neuronal contact onto the pericyte soma (p*) are indicated by *green* tint. L) Neuronal contacts broken down by vascular element and cell class.

Volume reconstructions in this dataset (Figure 3E-I) showed that patches of neuronal contact onto pericytes (green) and endothelial cells (yellow) could exhibit a wide range of sizes, from a single localized contact to larger patches comprising contacts from more than a dozen neuronal processes in a tight fascicle (Figure 3G,J). Some neuronal contacts were found on the peripheral pericyte processes (e.g., Figure 3F), but most were concentrated at or near the soma (Figure 3H). In this region, numerous spine-like appendages protruded from the pericyte’s surface, greatly increasing the available surface area for neuronal contacts (Figure 3F,K). Because these contacts may contribute to neurovascular coupling, we scanned them for any signs of presynaptic specialization. Though small features resembling synaptic vesicles could be found in some neurons near their vascular contacts, they did not concentrate near the contact, nor were there any other obvious structural features associated with conventional synaptic signaling (e.g. ribbons).

To learn whether fenestration of the glial sheath is unique to the capillaries of the IVP, we made similar reconstructions in the Helmstaedter dataset (Helmstaedter et al., 2013), which, though smaller than the Ding volume, includes all three vascular plexuses. It was prepared by a method that minimized intracellular detail and enhanced the extracellular spaces to simplify segmentation (Briggman et al., 2011). Conveniently for our study, the fixation and staining protocol enhanced visualization of the basement lamina surrounding endothelial cells and pericytes, rendering it nearly as dark as any structure in the block.

In this volume, the SVP was covered virtually completely by Müller and astrocytic glial processes, consistent with a recent report (Albargothy et al., 2023). However, we did identify two small gaps in the glial sheath where the SVP vessel was contacted by neurons ((Figures 4A, B-C *top* and S4A). In both cases, the contacts came from RGC axons, as revealed by skeleton reconstructions. Such gaps were much more common in the IVP (n=245) than in the DVP (n=17).

**Figure 4.**
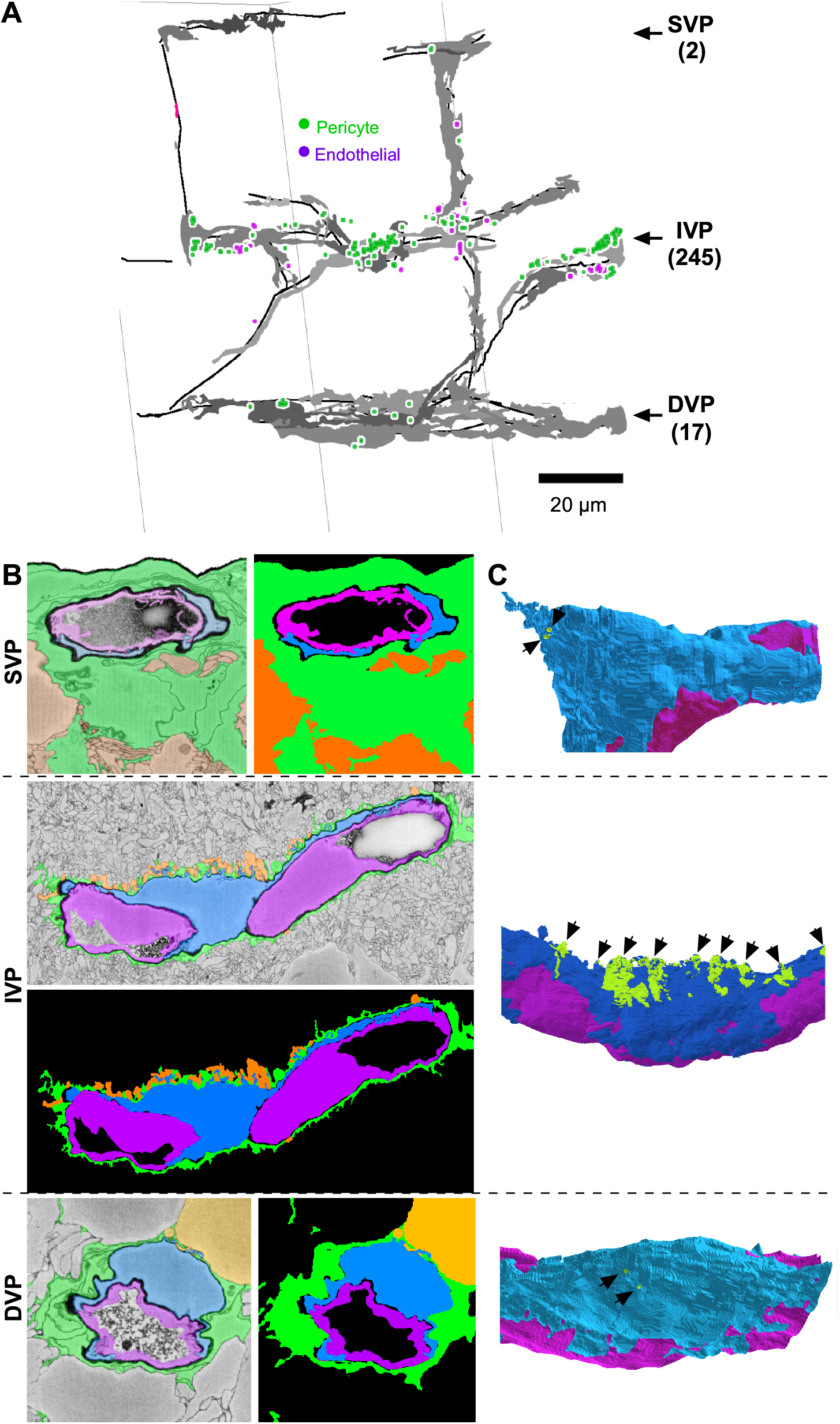
The IVP is more fenestrated than the SVP and DVP. A) Capillaries (black lines) and pericytes (gray volume) mapped for all three vascular layers in a SBFSEM dataset *(Helmstaedter et al*., *2013). Green* markers indicate a Müller ensheathment gap over a pericyte; *magenta* markers indicate ensheathment gaps exposing the vessel endothelium. The number of gaps in each layer is indicated in parentheses. B) Examples of near-complete Müller ensheathment in each vascular layer: Müller cells (green), pericytes (blue), endothelial cells (magenta). C) Partial 3D reconstructions of the capillaries and pericytes shown in B. Gaps in Müller sheath are green. SVP (*top*) and DVP (*bottom*) sheaths contain only two tiny gaps over pericyte somas, whereas some pericytes in the IVP have much larger gaps in Müller ensheathment. Exposed IVP pericyte somas often extend spines into the IPL (black arrows). See also Figures S3 and S4.

As in the Ding volume, many of the gaps were associated with spiny protrusions from the pericytes, concentrated perisomatically (Figure 4C *middle*). The basement lamina thinned dramatically around these spines (Figure S3A-B). We occasionally saw similar spine-like appendages in endothelial cells, again with thinned basement lamina and neuronal contacts. The lack of intracellular detail in this volume precluded assessment of possible synapse-like specializations.

To assess which cell types make direct contact with pericytes and endothelial cells in the IVP, we made skeleton reconstructions of the neurons that filled the gaps between Müller glial processes in the Ding volume. Some of these neurons were only partially contained within the volume of the dataset (e.g. wide-field amacrine and ganglion cells, and neurons near the block’s edges). These partial reconstructions typically provided enough information to determine the cell class (bipolar, amacrine, RGC) but were insufficient to identify cell types within a class.

All three cell classes with neurites near the IVP – ganglion cells, amacrine cells and bipolar cells – made direct contact onto pericytes and endothelial cells. Cell classes did not differ much in their proportion of contacts onto endothelial cells or pericytes (Figure 3L). Moreover, each cell class was represented by a large number of distinct types, together comprising virtually all of the neuronal types with processes stratifying near the IVP (Helmstaedter et al., 2013). For example, among amacrine cells familiar types such as the A17, VIP, and wide-field dopaminergic amacrine cells all made direct capillary contacts. Ganglion cells making numerous contacts onto both pericytes and endothelial cells included the F-mini Off, JamB, suppressed-by-contrast, and OFF sustained alpha types. Bipolar contacts were made mainly by OFF bipolar cone cell types with axonal arbors closest to the IVP, particularly types 1 and 2. Among all these cells, only the OFF bipolar cells exhibited an obvious bias towards pericytes.

### Fenestration of the glial sheath also permits neuron-to-vessel contacts in neocortex

To determine whether other CNS capillaries might exhibit similar fenestration of the glial sheath, we analyzed capillaries in two publicly available SBFSEM datasets of mouse somatic sensory cortex (Gour et al., 2021; Motta et al., 2019). In both samples, we found many instances where neuronal profiles penetrated through the astrocytic sheath to contact endothelial cells as well as pericytes. In the Gour dataset, obtained from an early postnatal mouse (P5; layers 2-4), pericyte contacts were most extensive near the pericyte soma, but were scattered widely over the cell’s surface and included distal processes (Figure S4). Many neuronal contacts were made onto endothelial spines. These could be short and stubby, or long and straight. Some spines penetrated through the glial sheath to contact neurons; these were clustered close to intercellular junctions between endothelial cells and the most distal processes of the pericytes (Figure S4G-K). Additional endothelial spines sprouted from the endothelial cell soma. One particularly long spine was almost completely covered by neuronal contacts (Figure S4G,HJ). Reconstructions of the contacting neurons showed that they were diverse, including dendritic spines, axonal varicosities, and some neuronal somas (Figure S4J,K). These findings were confirmed in the Motta volume, drawn from layer 4 of the same cortical region, but from a more mature mouse (P28). The automated segmentation publicly available on webknossos.org allowed us to quickly confirm that neuronal contacts onto both pericytes and endothelial cells come from diverse neuronal types.

### ATP-evoked Ca^2+^ activity in Müller sheaths

Ca^2+^ signals in Müller glia are known to play an important role in the vasodilation of IVP capillaries (Biesecker et al., 2016), but little is known about Ca^2+^ activity within the capillary sheaths. We labeled blood vessels with sulforhodamine (SR101) and loaded Müller cells with the fluorescent Ca^2+^ indicator Fluo-4 AM (see Star Methods) to monitor Müller cell Ca^2+^ responses to brief puffs of ATP (Figure 5). Consistent with previous work showing that Müller glia are sensitive to extracellular ATP (Liu & Wakakura, 1998; Newman & Zahs, 1997), we found that a brief puff of 5 mM ATP onto the retina’s inner limiting membrane elicited indicator signals in Müller cells at all retinal depths, including the sheaths surrounding capillaries in the IVP and DVP (Figure 5B-C, see Star Methods). We quantified these signals in two ways: 1) a gross measurement of Müller Fluo-4 signal in the vicinity of capillaries (i.e. ΔF/F of an ROI extending ∼10 µm in each direction from the capillary center) to account for loading or responsivity issues (Figure 5E), and 2) a normalized measure of sheath Ca^2+^ signaling and morphology relative to the capillary (ΔF z-score, Figure 5F; see Star Methods).

**Figure 5.**
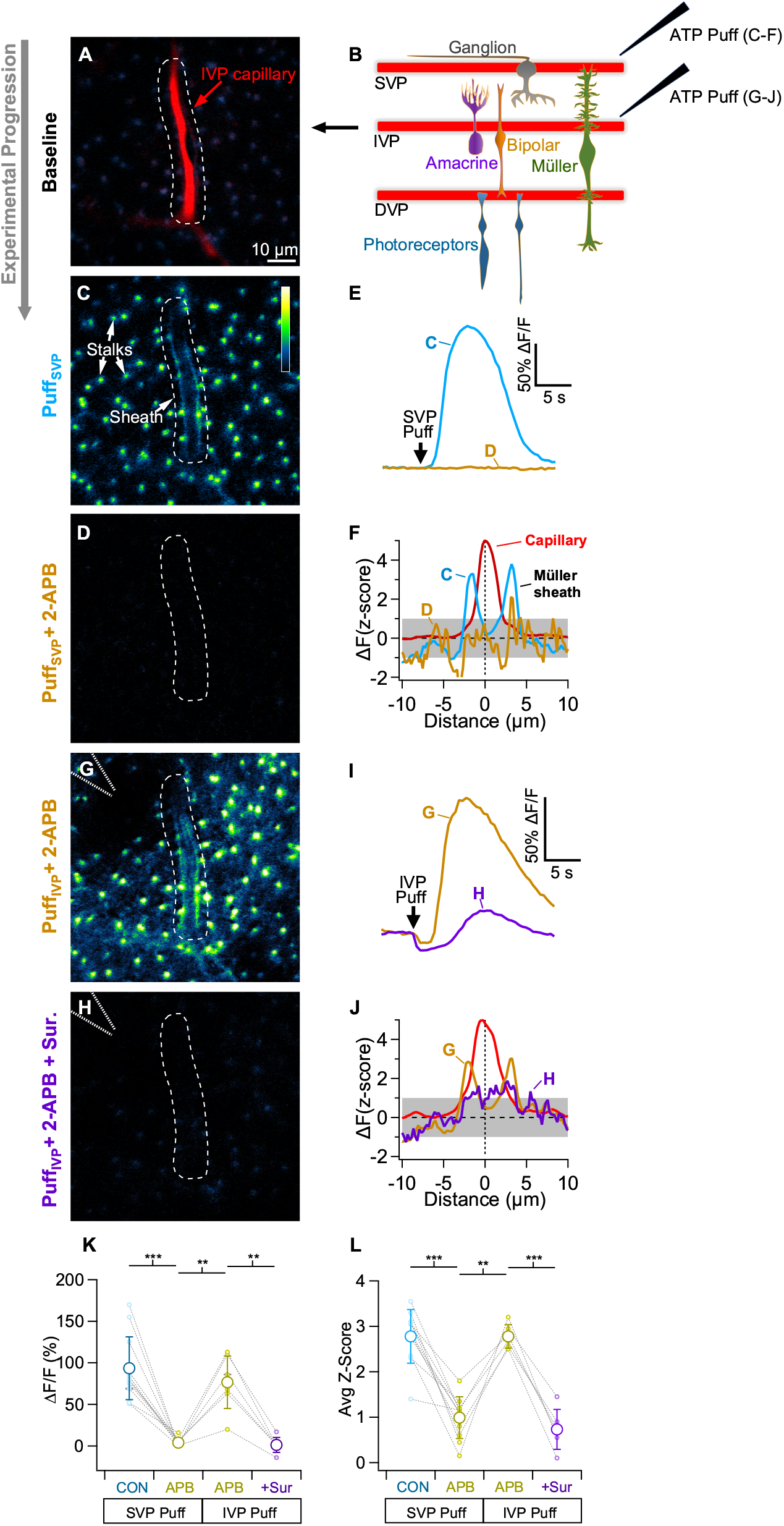
Purinergic Ca^2+^ signaling in Müller sheaths. A-B) Ca^2+^ (Fluo-4) signals in peri-capillary Müller glial processes evoked by ATP application to the vitreal retinal surface (just above the SVP). A) IVP capillary (filled with SR-101, *red*) and resting Ca^2+^ signals (Fluo-4, *blue-yellow*) in surrounding Müller glia. B) 5 mM ATP was puffed onto the (superficial) surface of the retina with a glass electrode (panels C-F). The electrode was then advanced into the IPL and ATP was puffed directly onto an IVP capillary (panels G-J). C) Baseline-subtracted Fluo-4 signals evoked by ATP puff on retina surface. D) As in C but in the presence of 100 μM 2-APB, an IP_3_R antagonist. Fluo-4 signal scale applies to all image panels. E) ATP-evoked ΔF/F responses for the ROIs in C and D. F) ATP-evoked fluorescence profiles of capillary sheaths in control and in 2-APB. Capillary signal is normalized to a z-score=5. G) Baseline-subtracted Ca^2+^ response to ATP puffed directly onto an IVP capillary in the presence of 2-APB. H) As in G, but with the addition of the purinergic receptor antagonist suramin. I) ATP-evoked ΔF/F responses for the ROIs in G and H. J) Fluorescence response profiles for direct capillary puffs in 2-APB and with the addition of suramin. K-L) Population data showing ΔF/F responses (K) and average z-score (L) of each vessel under each experimental condition.

Bath application of the IP_3_R antagonist 2-APB reduced the Fluo-4 responses of Müller sheaths to a puff of ATP at the retinal surface (ΔF/F_2-APB_=6±9% of control, z-score_2-APB_=38±21% of control, n=11 capillaries, Figure 5C-F). When the puff electrode was moved into the IPL, application of ATP directly onto the intermediate capillaries elicited robust Fluo-4 responses in ensheathing Müller processes, even in the presence of 2-APB (z-score_2-APB_: superficial puff=0.95±0.52 vs intermediate puff=2.7±0.2, n=6, Figure 5G,I,K). These responses were blocked by the purinergic receptor antagonist suramin (z-score=26±13% of 2-APB alone, ΔF/F=-8±26% of 2-APB alone, n=6, Figure 5H-L). These results indicate that the Müller processes that ensheathe capillaries express purinergic receptors and exhibit Ca^2+^ responses to ATP that are amplified and propagated by IP_3_R-mediated release from intracellular stores.

### Disruption of sheath activity in a mouse model of retinal degeneration

Many retinal diseases, including retinitis pigmentosa and diabetic retinopathy, trigger gliosis (Bringmann et al., 2006). To examine how these pathological processes affect Ca^2+^ signaling in the Müller sheaths and their association with retinal capillaries, we repeated our imaging experiments in the *rd10* mouse model of retinitis pigmentosa. *rd10* mice have a missense protein mutation in rod photoreceptors that leads to their early death; as rods die, gliosis is triggered and the retina begins to remodel (Bringmann et al., 2006). Wildtype (WT) and *rd10* retinas were loaded with Fluo-4 AM and SR101 and imaged while delivering ATP puffs on the retinal surface. This stimulus evoked robust indicator signals in Müller processes in the IVP in both WT (z-score=2.9±0.5, n=19) and *rd10* mice (z-score=3.1±0.6, n=21) (Figure 6A-C). On the other hand, in the DVP near rod synaptic terminals, we detected clear differences in the morphological relationship between Müller cells and capillaries. In WT mice, capillaries in the DVP were surrounded closely by a thin rim of Ca^2+^ activity (z-score = 3± 0.3, n=12, Figure 6D-F). In *rd10* mice, however, Müller processes in the deep capillary layer were more diffuse and irregular.

**Figure 6.**
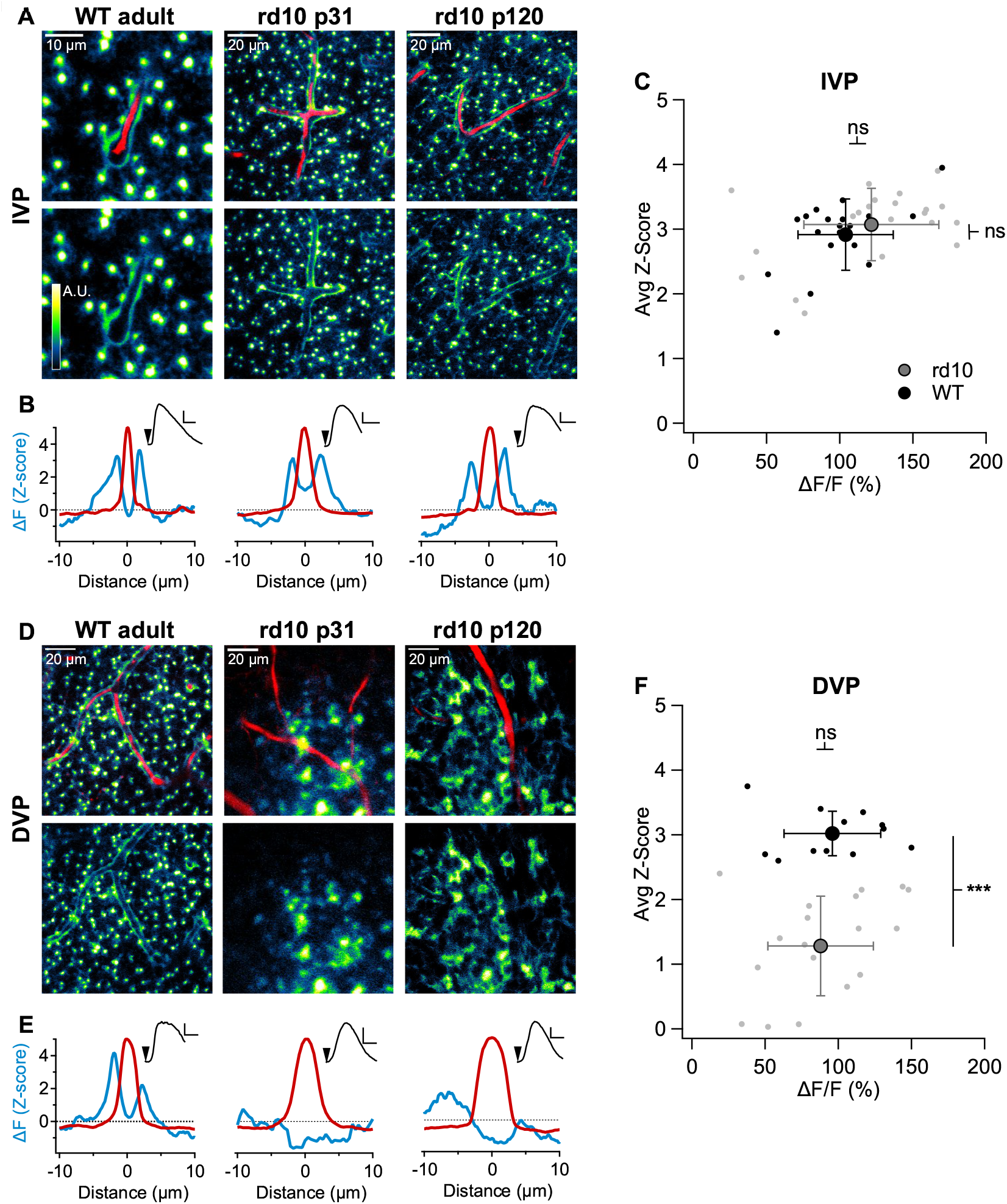
Müller sheaths are perturbed during retinal degeneration. A) Baseline subtracted image showing Müller cell Fluo-4 ΔF signals in the IVP in response to a superficial ATP puff. Capillaries are labeled with SR101. *left* WT adult, *middle* p31 *rd10, right* p121 *rd10*. B) ΔF Z-score (blue) and capillary profile (*red*, normalized to 5) vs. distance from the center of the capillary. *insets* ΔF/F Fluo-4 signals evoked by brief ATP puffs. C) Population data for sheath activity in the IVP. Each point represents one vessel segment; ≥1 mm of capillary was analyzed for each condition. D) Baseline subtracted image showing Müller activity in the DVP in response to a superficial ATP puff. E) As in B but for DVP capillaries. F) Population data for DVP sheath activity.

**Figure 7.**
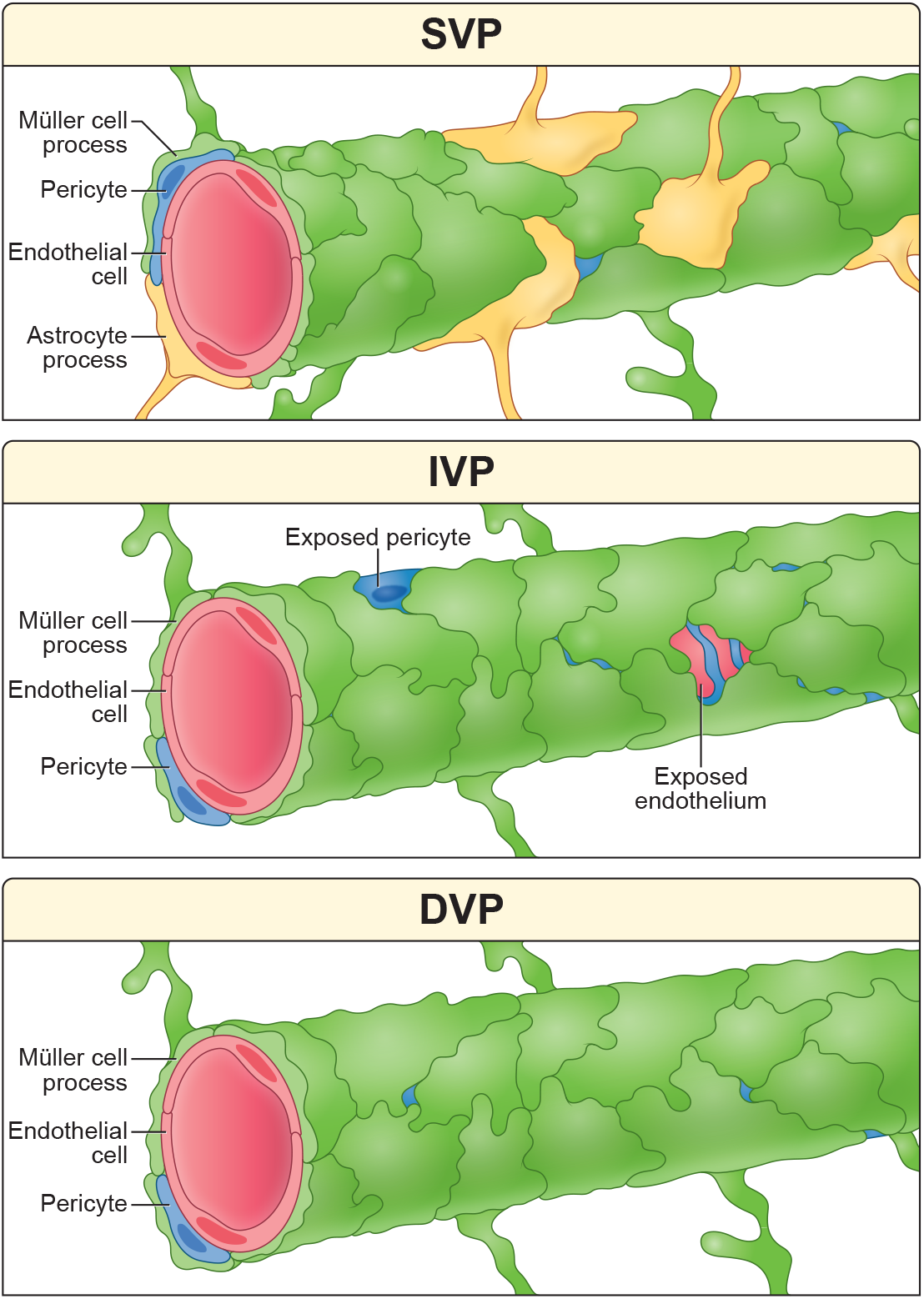
Illustrated conclusions from this study. *Top*, SVP capillaries are wrapped by astrocytes and Müller cells. Sheaths are essentially complete, limiting opportunities for neuronal contact. *Middle*, Müller glia cover IVP capillaries with fenestrated sheaths that allow neurons to directly contact endothelial cells and pericytes. *Bottom*, DVP capillaries are ensheathed by Müller glia with few gaps.

Although *rd10* Müller glia took up Fluo-4 and responded to ATP puffs (ΔF/F=88±36%), these indicator signals were no longer tightly associated with the vessels (z-score=1.3±0.8, n=19, Figure 6D-F). These results indicate that photoreceptor degeneration disrupts Müller sheaths in a layer-specific manner.

## Discussion

The retina’s vasculature supports the metabolic demands of visual processing. Here, we show that Müller glia encase capillaries in all three vascular layers of the mouse retina. Though these sheaths are nearly complete, gaps exist, especially in the IVP. When gliosis is triggered by photoreceptor degeneration, the Müller sheath is disrupted primarily in the DVP.

### Neuronal interactions with the perivascular elements

The near-complete (>90%) Müller glial coverage that we observe around retinal capillaries parallels capillary anatomy in the hippocampus, where the glial sheath is formed by astrocytic endfeet (Mathiisen et al., 2010), and in our preliminary analysis of the neocortex (Figure S4). Müller processes forming the sheath appear to adhere tightly to one another (Figure 2i), suggesting that glial sheaths insulate vessels from the surrounding milieu, permitting interactions between neurons and vessels only at sparsely distributed gaps. Neurons contact both endothelial cells and pericytes, apparently favoring pericyte somata. The contacting neurons appear to reflect the cell type diversity in the vicinity of the gap. Interestingly, the balance and ultrastructure of contacts made onto endothelial cells or pericytes appears to vary in different brain regions. For example, hippocampal endothelial cells are almost entirely wrapped by glia, with only a few small gaps allowing contacts with presumed microglial processes, while pericytes get more extensive neuronal contacts. In the retina and somatosensory cortex, we observed a more equal balance of endothelial and pericyte contacts. In the neocortex, we found many neuronal contacts at tight junctions between endothelial cells that contribute to the blood-brain barrier. In the IVP of the retina, many neuronal contacts were made onto pericyte somatic spines. These were far less common in the cortical volumes we studied, though at least one reconstructed endothelial cell extended a surprisingly long spine into the neuropil, where it was covered in neuronal contacts. In embryonic neocortex endothelial cells exhibit striking spinous appendages, some penetrating processes of neighboring neural progenitor cells (Wilsch-Brauninger et al., 2024).

The functional importance of these diverse neurovascular contacts remains unclear.

Physiological results suggest that Müller glia, like astrocytes, play an intermediary role in neurovascular coupling (Biesecker et al., 2016). Is there a role for the direct neural contacts reported here? These contacts are most prevalent in the IVP, which exhibits the largest activity-dependent changes in vessel diameter (Kornfield & Newman, 2014). Neurovascular contacts are made by diverse types of amacrine cells, ganglion cells and bipolar cells extending processes at the level of the IVP. These contacts lacked obvious presynaptic specializations, suggesting that these neurons may not use vesicular synaptic transmission to communicate with perivascular elements. A precedent for direct neurovascular communication exists in the glutamatergic signaling to smooth muscle cells in neocortical arterioles (Zhang et al., 2024).

Other signaling mechanisms may be involved, including gap junctions, transporter-mediated signaling and membrane-permeant molecules like nitric oxide. It is also possible that signals pass in the opposite direction, from vessels to neurons and glia, along with glucose and oxygen.

Molecular cues in the bloodstream modify neural stem cell activity during development and adulthood: specifically, in mouse neocortex blood cues help convert a single layer of proliferating progenitor cell into multiple distinct neuronal layers (Noctor et al., 2001). Given that Müller cells act as radial glia late-stage progenitors under certain conditions, cues in the blood may guide cell differentiation after injury (Bernardos et al., 2007; Jorstad et al., 2017; Wohlschlegel et al., 2023).

We observed marked variability in glial coverage of capillaries across and within the different brain regions examined. In the mouse retina, neurovascular contacts were much more prevalent in the IVP, though this requires confirmation in other volumes. Glial coverage in the cortex also appears to vary during development: Many contacts were observed in P5 mouse somatosensory cortex (Gour), with fewer seen at P28 (Motta). A survey of other cortical datasets (webknossos.org; MiCRONS project primary visual cortex) revealed many capillary segments with few if any neuronal contacts, similar to the virtually complete ensheathment of hippocampal endothelial cells (Mathiisen et al., 2010) and our reconstruction of a short stretch of IVP capillary from macaque retina. It is unclear what accounts for this diversity in neurovascular contacts – species, age, laminar location, and vessel branch order may all be factors.

### Müller sheaths and disease

Many neural diseases trigger gliosis, i.e., glial activation and/or proliferation. Here we examined Müller activity in the IVP and DVP in the *rd10* mouse model of retinal degeneration. Previous work in *rd10* mice showed that, as rod photoreceptors die, the blood-retinal barrier (BRB) breaks down first in the DVP, followed by the IVP and eventually the SVP (Ivanova et al., 2019). Our results from *rd10* mice (p31-p121) showing layer-specific alterations in Müller sheaths is consistent with this study, suggesting that barrier disruption coincides with loss of Müller ensheathment. Since degeneration in the *rd10* mouse model originates within rod photoreceptors, Müller processes in the deep layers may sense cell death and alter their function, as shown in P23H mice, another model of retinitis pigmentosa (Tomita et al., 2021). If Müller cells adapt to support dying photoreceptors, important resources might be diverted from BRB maintenance. Our data showing that sheath disruption, like BRB breakdown, is layer-specific suggest that alterations in Müller morphology might be highly compartmentalized. Future experiments may determine how Müller sheaths react in other diseases, such as diabetic retinopathy, where external factors in the blood (i.e., glucose) facilitate a global influence on the BRB.

## STAR METHODS

Detailed methods are provided in the online version of this paper and include the following:

- KEY RESOURCES TABLE
- RESOURCE AVAILABILITY
  - Lead contact
  - Materials availability
  - Data and code availability
- EXPERIMENTAL MODEL AND SUBJECT DETAILS
- METHOD DETAILS
  - Acquisition of macaque retinal SBFEM dataset
  - Analysis of SBFSEM data
  - Ca^2+^ imaging
  - Immunohistochemistry
- QUANTIFICATION AND STATISTICAL ANALYSIS

## KEY RESOURCES TABLE

| REAGENT or RESOURCE | SOURCE | IDENTIFIER |
| --- | --- | --- |
| *Antibodies* |  |  |
| GFAP (mouse polyclonal) | Sigma | G3893 |
| CD31 (goat polyclonal) | R&D | AF3628 |
| *Chemicals, peptides, and recombinant proteins* |  |  |
| Ames media | US Biological Life Sciences | A1372-25 |
| NaHCO<sub>3</sub> | Fischer Bioreagents | BP328-500 |
| Mg ATP | Millipore Sigma | A9187 |
| AlexaFluor 488 | ThermoFischer | A10436 |
| Sulforhodamine-101 | Millipore Sigma | S7635 |
| HEPES | Millipore Sigma | H3375 |
| Suramin | Millipore Sigma | S2671 |
| 2APB | Tocris Bioscience | 1224 |
| Triton X-100 | Millipore Sigma | 9036-19-5 |
| *Deposited data* |  |  |
| Igor files | TBD | TBD |
| *Experimental models: Organisms/strains* |  |  |
| mouse: wild-type: C57BL/6J | Jackson Laboratory | 000664 |
| mouse: Rd10 | Jackson Laboratory | 004297 |
| primate: Macaca nemestrina | Washington National Primate Center Tissue Distribution Program | n/a |
| *Software and algorithms* |  |  |
| Stage | Github | https://github.com/Stage-VSS/stage |
| Symphony | Github | https://github.com/Symphony-DAS/symphony-matlab |
| Data Analysis package | Github | https://github.com/Schwartz-AlaLaurila-Labs/sa-labs-extension |
| Zen | Zeiss | RRID: SCR_013672 |
| Igor Pro | Wavemetrics | https://www.wavemetrics.com/products/igorpro |
| Adobe Illustrator | Adobe | https://www.adobe.com/products/illustrator.html |
| Fiji | ImageJ | https://imagej.net/software/fiji/ |
| Knossos | Knossos | https://knossos.app/ |
| Paraview | Kitware | https://www.paraview.org/ |

## RESOURCE AVAILABILITY

### Lead Contact

Further information and requests for resources should be directed to and will be fulfilled by the lead contact, Jeffrey Diamond.

### Materials availability

This study did not generate new unique reagents.

### Data and code availability

- Original electrophysiology data have been deposited at TBD and are publicly available as of the date of publication. The DOI is listed in the key resources table. Microscopy data reported in this paper will be shared by the lead contact upon request.
- This paper does not report original code.
- Any additional information required to reanalyze the data reported in this paper is available from the lead contact upon request.

## EXPERIMENTAL MODEL AND SUBJECT DETAILS

We used wild type and rd10 mice of either sex for our experiments, ranging in age from p31-p121. Primate retinal tissue was obtained from an adult Macaca nemestrina (macaque) through the Washington National Primate Center Tissue Distribution Program. All procedures were approved by the University of Washington, University of Wisconsin-Madison and/or NINDS Institutional Animal Care and Use Committee (ASP-1344).

## METHOD DETAILS

### Acquisition of macaque retinal SBFEM dataset

The macaque volume was prepared by dissecting the living retina in bicarbonate buffered Ames solution. A piece was cut from peripheral retina and immersion fixed in 4% glutaraldehyde in 0.1 M sodium cacodylate buffer (pH 7.4). The fixed sample was rinsed in cacodylate buffer and processed for serial block face scanning electron microscopy following the protocol of (Della Santina et al., 2016). Sectioning and imaging were performed on a 3View (Zeiss) serial block face scanning electron microscope at the University of Wisconsin-Madison School of Medicine Electron microscopy facility. Images were acquired at 5nm/pixel XY resolution at a section thickness (Z) of 50nm. Multi-montage acquisition was used to capture a wide field of view across the retina with each tile spanning ∼45 µm. Sectioning and imaging was performed across the vertical cross-section.

### Analysis of SBFSEM data

Mouse retinal and cortical SBFEM datasets have been described in detail in the papers that introduced them and published work (Ding et al., 2016; Gour et al., 2021; Helmstaedter et al., 2013; Motta et al., 2019; Pallotto et al., 2015). All are publicly available through webknossos.org. The Ding volume measures ∼50 × 210 × 260 µm, with a voxel size of 13.2 × 13.2 × 26 nm (Ding et al., 2016). This resolution allowed for visualization of vesicles and chemical synapses but doesn’t permit identification of gap junctions. The Pallotto volume was 60 × 80 × 80 µm with a voxel size like that of the Ding block. The preparation of the Pallotto tissue was optimized to preserve extracellular space within the retinal tissue (Pallotto et al., 2015). The Helmstaedter block was approximately 132 x 114 x 80 µm (Helmstaedter et al., 2013). This tissue was processed in a manner that enhanced extracellular contrast and cleared intracellular organelles to facilitate crowd-sourced reconstructions. The fourth block consisted of standard EM preparation with non-human primate retina, with a volume of ∼90 x 90 x 90 µm. The Motta block was 61.8 x 94.8 x 92.6 µm, with a voxel size of 11.24 x 11.24 x 28 nm, and was taken from layer four of the somatosensory cortex (Motta et al., 2019). The Gour block was 76 x 92 x 78 μm, with a voxel size of 11 x 11 x 30 nm, and was taken from the neocortex (P5,(Gour et al., 2021)).

Müller glia stalks were identified using characteristic dense internal filament structures that are absent from other retinal cell types. Stalk locations were identified and used to perform mosaic analysis of the glial population. After stalk identification, other glial processes extending out from the stalk were be traced and reconstructed. Vasculature was readily identified at low magnification from the hollow profile and continuous ring of endothelial cells. Pericytes were identified by their close association with endothelial cells. Their somas and long flattened processes clung to the vessels, exhibiting characteristic peg-and-socket arrangements of their fine terminal processes with endothelial and glial cells.

The Pallotto and Ding datasets were reconstructed and annotated using the EM analysis software Knossos (https://knossos.app/). The Ding, Helmstaedter, and mouse cortical datasets were annotated using the webknossos.org interface. Skeleton and 3D volumetric reconstructions from these datasets are webknossos screenshots. For the macaque block, reconstructions and annotations were done using ImageJ (http://imagej.nih.gov), with renderings of skeletons and volumes generated with the visualization software Paraview (Ahrens et al., 2005).

### Ca^2+^ imaging

Prior to experiments mice were deeply anaesthetized with isoflurane (Baxter) and euthanized vis cervical dislocation. Once euthanized the animals were enucleated and the eyes were submerged in Ames medium (room temperature, continuously bubbled with 95% O2/5% CO2 gas). A small incision was made at the limbus, and scissors we then used to remove the cornea. The lens and vitreous were removed with forceps, and tissue was stored for up to 5 hours in this condition. Individual pieces of retina were isolated from the pigment epithelium and placed on a 10mm coverslip coated with poly-l-lysine as needed. The edge of a kim wipe was used to remove excess solution prior to application of 200 µL of freshly gassed Ames, containing 62.5 µM Fluo-4 AM and 1 mM SR101, directly to the top of the wicked-down retina. The coverslip was mounted in a recording chamber and placed in a dark incubator with continuous carbogen flow for 30 min. After incubation the recording chamber was placed under the microscope and superfused with freshly gassed Ames (∼8 mL/min) for at least 10 minutes before collecting data. Ca^2+^ signals and blood vessels were imaged with a Zeiss LSM510 confocal microscope (λ=488 nm). A 8-10 MΩ glass electrode filled with HEPES-buffered Ames containing 5mM ATP was used to puff (Picospritzer, 5 psi, 0.05-1 s in duration) ATP on the retinal surface (i.e. ILM) or directly onto the intermediate capillaries.

### Immunohistochemistry

Dissected Retina was post-fixed containing 4% Paraformaldehyde (PFA) in 1x PBS for 30 minutes at RT, rinsed with PBS and store in 4 °C until use. Stored retinas then were blocked for 24 hours in a PBS solution plus 0.5% Triton X-100 and containing 10% normal donkey serum (NDS). Primary antibodies were diluted in the same solution and applied for 72-96 hours, followed by incubation for 24 h in the appropriate secondary antibodies. We used GFAP (Sigma, 1:1000) and CD31 (R&D, 1:1000) to label astrocytes and blood vessels, respectively. All steps were completed at in 4 °C.

After staining, the tissue was flat-mounted on a slide, ganglion cell layer up, and cover slipped using Vectashield mounting medium (H-1000, Vector Laboratories). Immunoreactivity was visualized and acquired with a confocal microscope (Zeiss LSM780) using 20x air or 63x/1.4 oil objectives. The selected images were cropped and aligned with Fiji (https://imagej.net/software/fiji/).

## QUANTIFICATION AND STATISTICAL ANALYSIS

For Ca^2+^ imaging experiments, time series were analyzed in Fiji, and group data was compiled in Igor pro. We focused on capillaries in the intermediate and deep layers, and only analyzed sections that were fully within the plane of focus. Brief ATP puffs evoked radially expanding waves of Ca^2+^ activity, and analysis was also limited to within these regions. From these movies two measurements were extracted: 1) DF/F for a ROI that extended approximately three vessel diameters in both directions from the center of the capillary of interest—as a general readout of puff-evoked Ca^2+^ signals in Müller glia, and 2) a vessel-centric 2D analysis of the structure of local Ca^2+^ signals. For the vessel-centric 2D analysis 8-10 movies frames were averaged prior to stimulation and at the peak of the puff-evoked Ca^2+^ signal. The averaged baseline image was then subtracted from the averaged stimulation image to isolate the puff-evoked Ca^2+^ response. Ca^2+^ signals attributed to Müller stalks were then digitally removed from the baseline-subtracted image by: 1) thresholding the top 2-4% of pixel values of the averaged peak image (i.e. not baseline subtracted), 2) converting the thresholded pixels into a mask, 3) dilating the binary mask two-three times and 4) applying the mask to the baseline-subtracted image. This process converted stalk pixels to NaN which were then excluded from analysis. The resultant image was then combined with the SR101 labelling of blood vessels. Capillaries with curvature in the plane of focus were straightened with Fiji’s *straighten* function before taking vessel-centric measurements. To quantify the strength and locality of puff-evoked sheath activity we examined fluorescence signals as they extend from the center of the vessel. Average fluorescence signals were collapsed along the vessel axis for the entire length of the in-focus capillary of interest, and to a minimum of 20 µm on either side of the vessel. These profiles were converted to z-scores, and we quantified the average peak fluorescence from both sides of the vessel. Fluorescence originating from the blood vessels was normalized to a z-score of 5 for easy viewing.

All statistical analysis was performed using Igor Pro. Reported n values indicate number of cells. Normally distributed (as determined by the Jarque-Bera test) group data values are reported as mean ± SD and compared with a *t*-test. Significant differences were concluded if *p* < .05.

## SUPPLEMENTAL INFORMATION

Document S1. Figures S1-S4.

## ACKNOWLEDGEMENTS

This work was supported by the NINDS Intramural Research Program (NS003145 to JSD), UW2020 WARF discovery award (MH), and NIH-NEI R01EY12793 (DMB).

