## Supplemental Data for "The retina’s neurovascular unit: Müller glial sheaths and neuronal contacts"

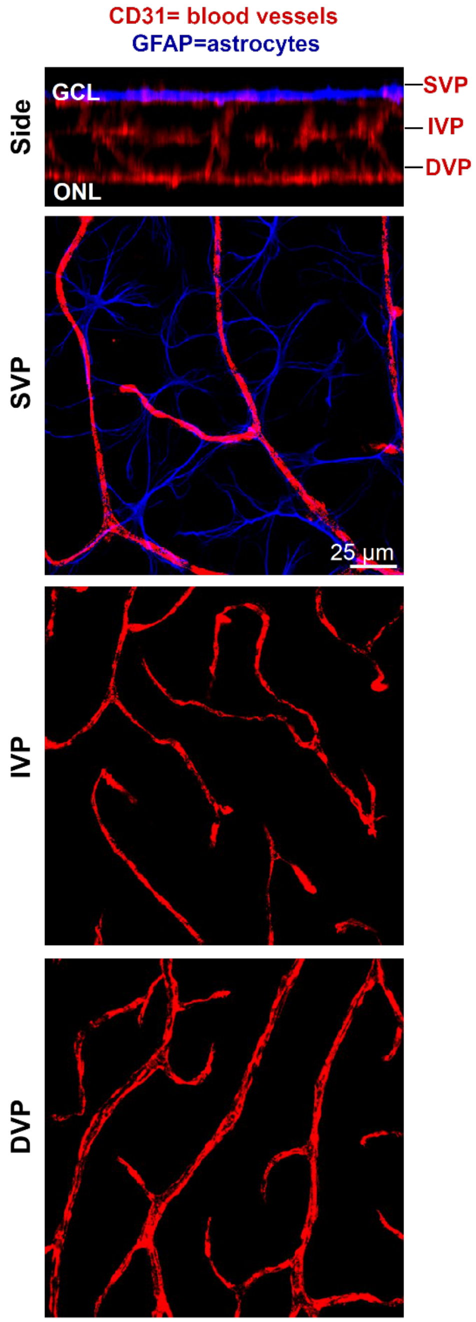


**Figure S1 (related to Figure 1).** *Retinal astrocytes are confined to the superficial vascular layer.* Immunohistochemistry labeling of retinal blood vessels (red, CD31) and astrocytes (blue, GFAP).

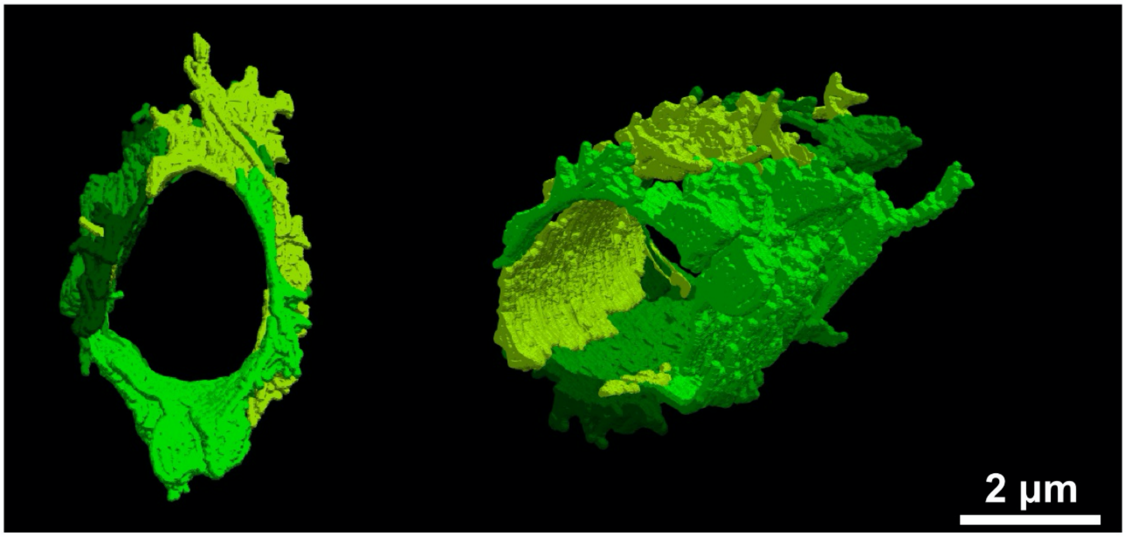


**Figure S2 (related to Figure 2).** *Recolored images from Figure 2E according to Müller parent.* Processes in Figure 2E were traced back to the parent Müller cell. Each shade of green represents processes from a common Müller parent.

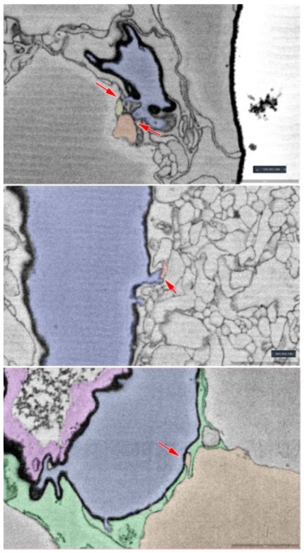

**Figure S3 (related to Figure 4).** *Sites of contact between neurons and pericytes (blue tint) in all three vascular layers as viewed in single EM planes from the Helmstaedter volume. top* Contact of SVP pericyte protrusions with two RGC axons (yellow, orange). *middle* Spine (red arrow) of an IVP pericyte soma contacting a fine neuronal process (pink). Note the thinning of the basement lamina around the spine. *bottom* Protrusion from a bipolar cell soma contacts a pericyte soma (red arrow). In this panel, green tint marks Müller cell processes and pink marks endothelium.

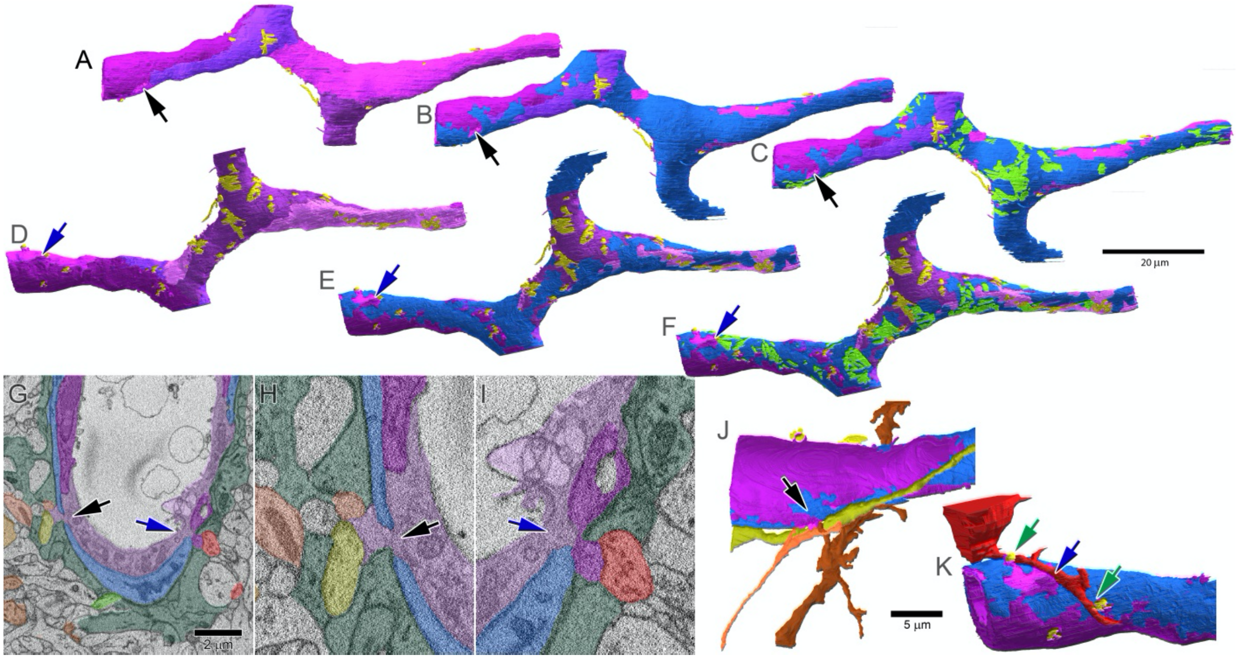


**Figure S4 (related to Figure 4).** *Reconstructions of neuronal contacts onto a neocortical capillary using a publicly available SBFSEM dataset of mouse somatosensory cortex* (P5; (Gour et al., 2021)). Endothelial cells are *purple* and pericytes are *blue* in all panels. A-F: Six views of the capillary, highlighting different structural features. Top row (A-C) show the vessel from one perspective and the bottom row (D-F) after rotation of the capillary around its long axis to reveal the reverse side. Left pair vessel reconstructions (A,D) show endothelial cells and their neuronal contacts (*yellow*). Individual endothelial cells are shown in slightly different shades to reveal intercellular junctions. Middle pair of views (B,E): same as A,D but with the addition of the pericyte (*blue*). Right views (C,F): same but with the addition of neuronal contacts to the pericyte (*green*). The huge patch of neuronal contacts in C marks the location of the pericyte's soma. An endothelial soma appears in D (center) marked by large patches of neuronal contact (yellow). Note that neuron-to-endothelial contacts are concentrated on one side of the capillary (D-F) and (except near the soma) cluster near distal pericyte processes. They are also concentrated at intercellular junctions between endothelial cells, especially on short endothelial spines, but also on long somatic spines (E; middle). G-I: electron micrograph drawn from a single plane of the dataset passing through the same capillary (coordinates: 1680, 832, 538); with selected cellular profiles tinted to highlight features of interest. Endothelium: *purple*; pericyte: *blue*; astrocytes forming vascular sheath: *dark green*; neurons contacting endothelium: *warm colors*; neuron contacting pericyte: *green*. *Black arrow* in G and in the expanded view in H marks the site of a short endothelial spine that skirts the edge of the pericyte and penetrates through the otherwise continuous astrocytic sheath (*dark green*) into the parenchema, where it receives direct contacts from several neuronal processes. This location is also marked by black arrows in the expanded view in H as well in the 3D reconstructions of A-C and J. *Blue arrow* in C (as well as D-F and K) marks the site of a similar endothelial spine with neuronal contacts; this spine derives from the other endothelial cell forming the capillary wall here. Both endothelial spines lie near the boundary between the two endothelials cells (two shades of purple), as well as near distal pericyte processes that receive local neuronal contact (C and F; arrows). J,K: local volumetric reconstructions of the neuronal processes making contact at the sites marked in other panels by the black arrow (J) and blue arrow (K). These include presumed axons (*orange* and *olive* in J) as well as spiny dendrites (*dark orange* in J; proximal dendrite in *red* in K, with partial somatic reconstruction) which makes two additional contacts onto the endothelium (*green arrows*), one of them again at an intercellular junction between endothelial cells (right green arrow).
